# Fatty acid metabolic interactome atlas linked to cellular longevity

**DOI:** 10.64898/2026.05.20.726705

**Authors:** Arshia Naaz, Mingtong Gao, Yizhong Zhang, Rajkumar Dorajoo, Brian K. Kennedy, Mohammad Alfatah

**Author notes:** To whom the correspondence should be addressed. (Mohammad Alfatah) (Arshia Naaz).

## Abstract

Fatty acid biosynthesis is a central metabolic process required for membrane formation, organelle maintenance, and cellular proliferation, yet its broader relationship with stress responses and cellular aging remains incompletely understood. Here, we combined human and yeast interactome analyses with transcriptomic profiling and chronological lifespan assays to investigate the systems-level organization of fatty acid metabolic pathways and their relationship to cellular longevity. Integrated interactome mapping of mammalian fatty acid metabolic regulators, including *ACACA, FASN, SCD, ACSL, ELOVL*, and *SLC27* family proteins, together with conserved yeast orthologs including *ACC1, FAS1/FAS2, ELO1-3, OLE1, FAA1-4, ACS1/ACS2*, and *FAT1* revealed strong enrichment of anabolic growth regulation, membrane organization, vesicle trafficking, proteostasis, and endoplasmic reticulum (ER)-associated stress pathways. In yeast, fatty acid metabolic networks segregated into distinct anabolic and membrane-associated functional modules. Pharmacological inhibition of fatty acid synthesis using cerulenin suppressed cellular proliferation while extending chronological lifespan and induced broad downregulation of translation-associated anabolic pathways together with activation of stress-associated and membrane lipid remodeling programs. ER-associated interactome analyses of *DPAGT1* and *ALG7* further identified strong enrichment of membrane trafficking and unfolded protein response (UPR)-associated pathways, while pharmacological ER stress induction using tunicamycin also promoted enhanced chronological longevity. Collectively, our findings support a conserved model in which perturbation of fatty acid metabolic pathways remodels anabolic growth and ER-associated stress responses to promote cellular longevity.

## INTRODUCTION

Fatty acid biosynthesis is a fundamental cellular process required for membrane formation, organelle maintenance, energy storage, and cellular proliferation ^1–8^. In mammalian systems, fatty acid metabolic pathways involve coordinated regulation of fatty acid synthesis, elongation, desaturation, activation, and transport mediated by central metabolic regulators including acetyl-CoA carboxylase (*ACACA*), fatty acid synthase (*FASN*), stearoyl-CoA desaturases (*SCD*), acyl-CoA synthetase long-chain family members (*ACSL*), elongation of very long-chain fatty acid proteins (*ELOVL*), and solute carrier family transporters (*SLC27*) ^8–12^. In the genetically tractable unicellular yeast model *Saccharomyces cerevisiae*, conserved fatty acid metabolic pathways are regulated through orthologous enzymes including *ACC1, FAS1/FAS2, ELO1-3, OLE1, FAA1-4, ACS1/ACS2*, and *FAT1*, which collectively coordinate fatty acid synthesis, elongation, desaturation, activation, and membrane lipid remodeling ^13–19^.

Fatty acid metabolism is closely linked to central carbon metabolism, where glucose-derived acetyl-CoA serves as a major substrate for lipid biosynthesis and membrane formation. Fatty acid metabolic pathways further contribute to the synthesis and remodeling of complex membrane lipids, including phospholipids and sphingolipids, which are essential for organelle integrity, membrane trafficking, stress signaling, and cellular adaptation. Increasing evidence suggests that perturbations in lipid metabolic organization influence membrane homeostasis, endoplasmic reticulum (ER) function, mitochondrial activity, proteostasis, and oxidative stress responses ^8,9,20,21^. Consequently, alterations in fatty acid metabolism and membrane lipid homeostasis have increasingly been implicated in aging and age-associated diseases, including metabolic disorders, cardiovascular dysfunction, neurodegeneration, and cancer ^22–29^. However, despite increasing recognition of the importance of lipid metabolic regulation in aging biology, how fatty acid metabolic networks are functionally linked to anabolic growth control, ER-associated stress adaptation, and cellular longevity remains incompletely understood ^30–35^.

In the present study, we performed integrated interactome analyses to uncover the systems-level organization of fatty acid metabolic pathways in mammalian and yeast systems. Through integrated network analyses, we generated fatty acid metabolic interactome atlases and identified strong coupling between lipid metabolic pathways, anabolic growth-associated programs, membrane organization, proteostasis, and endoplasmic reticulum (ER)-associated stress pathways. Functional analyses of these conserved signatures further demonstrated that pharmacological inhibition of fatty acid synthesis using cerulenin suppresses anabolic growth programs while promoting enhanced chronological longevity in yeast. In parallel, pharmacological induction of ER stress using tunicamycin also extended chronological lifespan, supporting a functional link between ER-associated stress responses and cellular survival. Together, our findings support a conserved model in which perturbation of fatty acid metabolic pathways remodels anabolic growth and ER-associated stress responses to promote cellular longevity.

## RESULTS

### Human fatty acid metabolic interactome atlas reveals stress- and proteostasis-associated network organization

To systematically investigate the molecular organization of fatty acid metabolic pathways, we retrieved molecular interactors of major human fatty acid synthesis, elongation, desaturation, and transport regulators from the Alliance of Genome Resources database, including *ACACA, FASN, SCD, ACSL, ELOVL*, and *SLC27* family members **(Supplementary Data 1)**. The integrated interaction datasets were subsequently analyzed using Metascape to generate a human fatty acid metabolic interactome atlas **(Figures 1A–1I)** ^36^.

**Figure 1.**
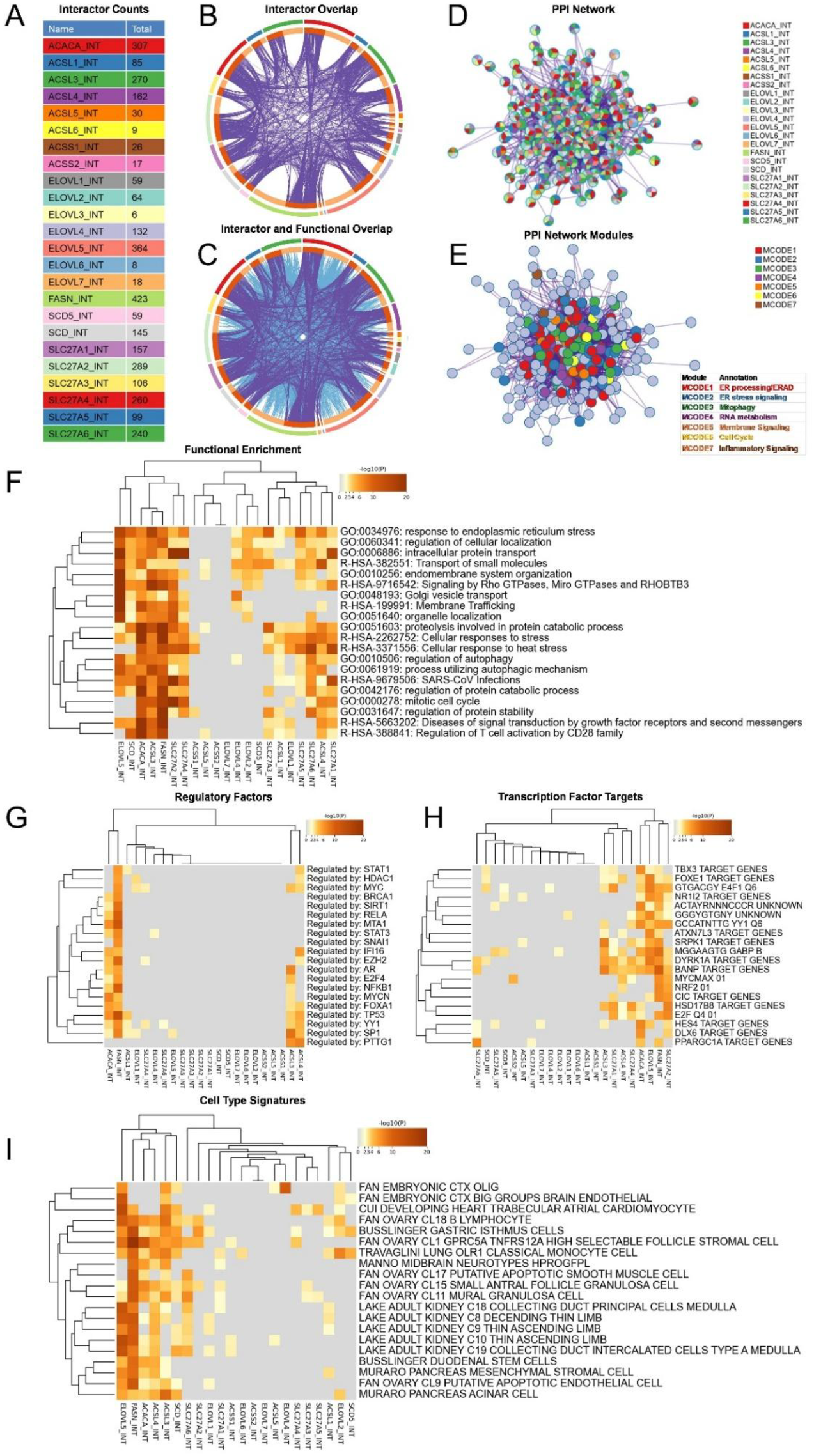
Integrated interactome analysis of human fatty acid metabolic regulators. (A) Total molecular interactors identified for human fatty acid metabolic regulators, including *ACACA, FASN, SCD, SCD5, ACSL1-6, ACSS1-2, ELOVL1-7*, and *SLC27A1-6* family proteins retrieved from the Alliance of Genome Resources database. Each fatty acid metabolic regulator is represented by a distinct color code maintained throughout the figure. (B) Circos plot showing overlap between molecular interactors among human fatty acid metabolic regulators. Outer color annotations correspond to individual regulators as shown in panel A. (C) Integrated interactome and functional overlap network illustrating shared molecular interactors and enriched biological pathways across fatty acid metabolic regulators. (D) Protein–protein interaction (PPI) network generated from integrated fatty acid metabolic interactors. Node colors indicate corresponding fatty acid metabolic regulators. (E) MCODE-based PPI network module analysis identifying enriched subnetworks associated with ER processing/ERAD, ER stress signaling, mitophagy, RNA metabolism, membrane signaling, cell cycle regulation, and inflammatory signaling pathways. (F) Heatmap showing functional enrichment analysis of Gene Ontology (GO) biological processes and pathway terms associated with individual fatty acid metabolic interactomes. (G) Regulatory factor enrichment analysis identifying transcriptional regulatory signatures associated with fatty acid metabolic interactomes. (H) Transcription factor target enrichment analysis showing conserved transcriptional programs linked to fatty acid metabolic pathways. (I) Cell-type signature enrichment analysis associated with human fatty acid metabolic interactomes. Color intensity in heatmaps represents enrichment significance (-log10 P value).

Interactor overlap analysis revealed substantial sharing of molecular interactors among multiple fatty acid metabolic regulators, indicating coordinated and interconnected network organization across fatty acid synthesis, elongation, desaturation, and transport pathways **(Figure 1B)**. Integration of both molecular interactors and functional ontology relationships further demonstrated broader overlap among fatty acid metabolic pathways and associated biological functions **(Figure 1C)**.

Protein–protein interaction (PPI) network analysis identified densely connected interaction networks among fatty acid metabolic regulators **(Figure 1D)**. Subsequent network module analysis revealed several functionally enriched subnetworks associated with proteostasis, ER stress signaling, mitophagy, RNA metabolism, membrane signaling, cell cycle progression, and inflammatory signaling **(Figure 1E)**.

Enriched ontology cluster analysis across integrated interactor studies demonstrated significant associations with endoplasmic reticulum stress responses, intracellular protein transport, membrane trafficking, endomembrane organization, autophagy, protein catabolism, cellular stress responses, and regulation of protein stability **(Figure 1F)**. Upstream transcriptional regulatory analysis identified signaling programs linked to *STAT1, MYC, BRCA1, RELA, TP53, SIRT1*, and *FOXA1* **(Figure 1G)**, while transcription factor target enrichment analysis highlighted additional metabolic and stress-associated regulatory signatures **(Figure 1H)**. Cell-type enrichment analysis further associated the fatty acid metabolic interactome with brain endothelial, cardiomyocyte, renal epithelial, pancreatic, immune-associated, ovarian stromal/granulosa, smooth muscle, and stem/progenitor-related cellular signatures **(Figure 1I)**, supporting broader links between lipid metabolic organization and vascular, metabolic, developmental, reproductive, and stress-responsive cellular systems. Collectively, these analyses indicate that human fatty acid metabolic pathways are integrated with conserved stress-responsive, proteostatic, membrane-associated, and nutrient-regulatory cellular programs.

### *FAS1/FAS2* interactome analysis reveals coupling of fatty acid synthesis with anabolic growth and stress-associated cellular pathways

Given the conserved association of mammalian fatty acid metabolic networks with stress-responsive and growth-regulatory pathways, we next investigated whether similar organizational features are present within the yeast fatty acid synthase complex composed of Fas1 and Fas2. To examine the broader cellular organization associated with fatty acid biosynthesis, we retrieved and integrated both physical and genetic interactors of *FAS1* and *FAS2* from the *Saccharomyces Genome Database* (SGD) **(Supplementary Data 2)** and subsequently performed Metascape analysis to construct a yeast fatty acid synthase interaction network ^36^.

Interaction network analysis revealed extensive overlap and connectivity between *FAS1* and *FAS2*, consistent with their coordinated function within the fatty acid synthase complex **(Figures 2A and 2B)**. Expansion of the *FAS1/FAS2* interaction landscape identified a broad range of associated proteins linked to RNA metabolism, protein homeostasis, cellular organization, and anabolic growth-associated pathways. Functional enrichment analysis further demonstrated strong enrichment for cytoplasmic ribosomal proteins, ribosome biogenesis, RNA processing, chromatin organization, transcription-associated pathways, proteasome assembly, protein localization, and regulation of catabolic processes **(Figures 2C and 2D)**. Multiple pathways associated with cellular stress responses, mitochondrial organization, autophagy-related regulation, heat stress responses, sphingolipid homeostasis, and endoplasmic reticulum-associated organization were also enriched within the *FAS1/FAS2* interactome.

**Figure 2.**
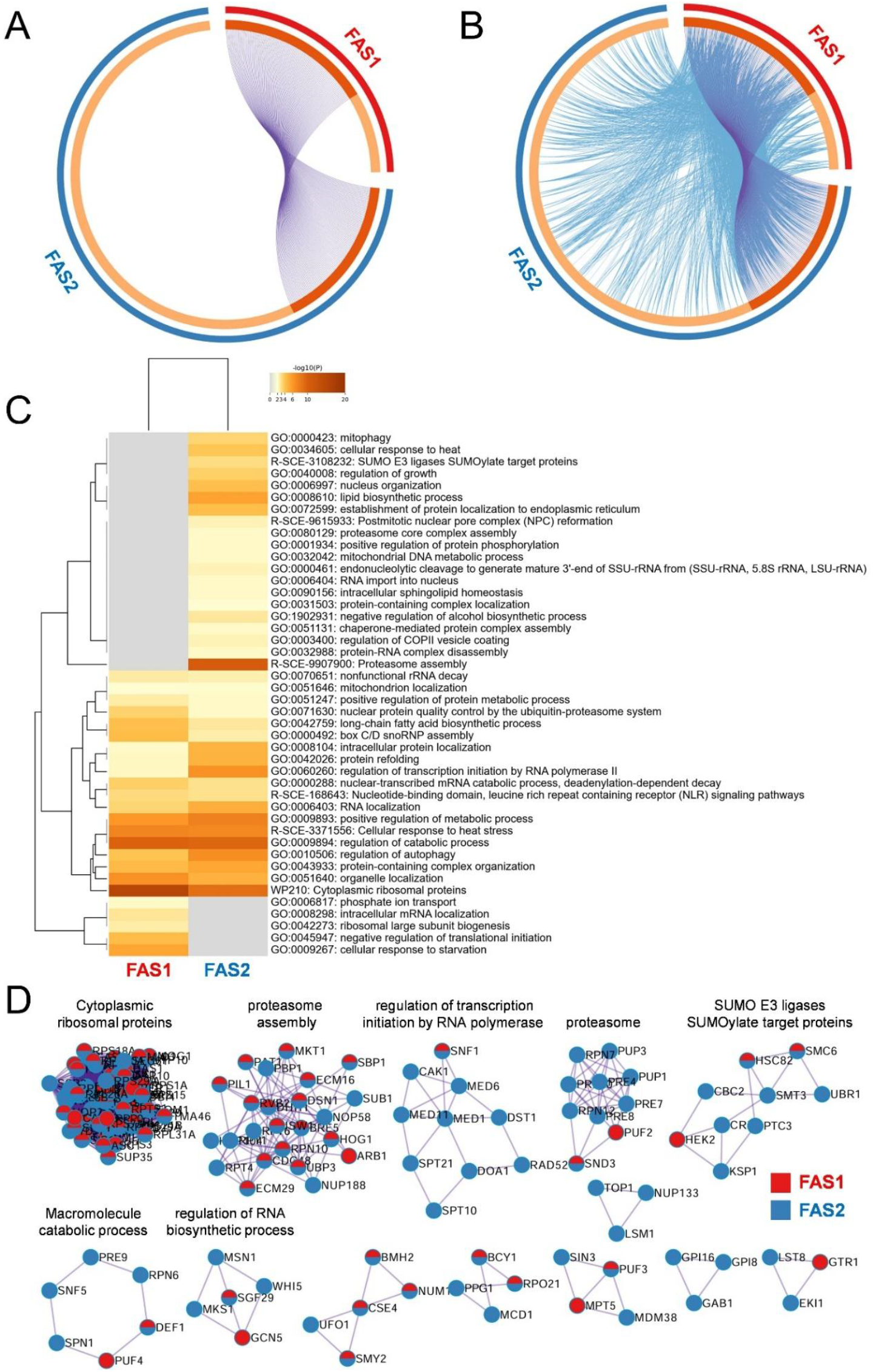
Integrated interactome analysis of the yeast fatty acid synthase complex *FAS1/FAS2*. (A) Circos plot showing overlap between physical and genetic interactors of *FAS1* and *FAS2* retrieved from the *Saccharomyces* Genome Database (SGD). Outer color annotations represent *FAS1* (red) and *FAS2* (blue) interactome networks. (B) Integrated interactome and functional overlap network illustrating shared molecular interactors and associated biological pathways between *FAS1* and *FAS2*. (C) Heatmap showing functional enrichment analysis of Gene Ontology (GO) biological processes and pathway terms associated with *FAS1*- and *FAS2*-associated interactomes. Color intensity represents enrichment significance (-log10 P value). (D) MCODE-based protein–protein interaction (PPI) network module analysis of the *FAS1/FAS2* interactome identifying representative subnetworks associated with cytoplasmic ribosomal proteins, proteasome assembly, transcription-associated pathways, SUMOylation-associated pathways, macromolecule catabolic processes, RNA biosynthetic regulation, and intracellular protein localization. Node colors indicate *FAS1*- and *FAS2*-associated interactors.

Although substantial overlap was observed between *FAS1*- and *FAS2*-associated networks, comparative enrichment analysis also revealed partially distinct organizational features. *FAS1*-associated interactors displayed relatively stronger enrichment for translation-associated pathways, ribosome biogenesis, mRNA localization, and starvation-responsive processes, whereas *FAS2*-associated networks showed comparatively stronger associations with proteostasis, protein refolding, intracellular protein localization, ER-associated organization, and stress-responsive pathways including heat stress adaptation and sphingolipid homeostasis **(Figures 2C and 2D)**. These findings suggest that, despite functioning together within the fatty acid synthase complex, *FAS1* and *FAS2* exhibit partially distinct connectivity patterns linking fatty acid biosynthesis with anabolic and stress-associated cellular programs.

### Yeast fatty acid metabolic interactome analysis reveals integration with anabolic growth and membrane-associated cellular pathways

To further investigate the systems-level organization of yeast fatty acid metabolic pathways beyond the Fas1/Fas2 synthase complex, we expanded the interactome analysis to include additional regulators involved in fatty acid biosynthesis, elongation, activation, desaturation, and lipid metabolism, including *ACC1, ACS1, ACS2, ELO1-3, FAA1-4, FAS1, FAS2, FAT1*, and *OLE1*. Physical and genetic interaction datasets were retrieved from the *Saccharomyces* Genome Database (SGD) **(Supplementary Data 2)** and analyzed using Metascape to construct an integrated yeast fatty acid metabolic interactome atlas **(Figures 3A–3G)** ^36^.

**Figure 3.**
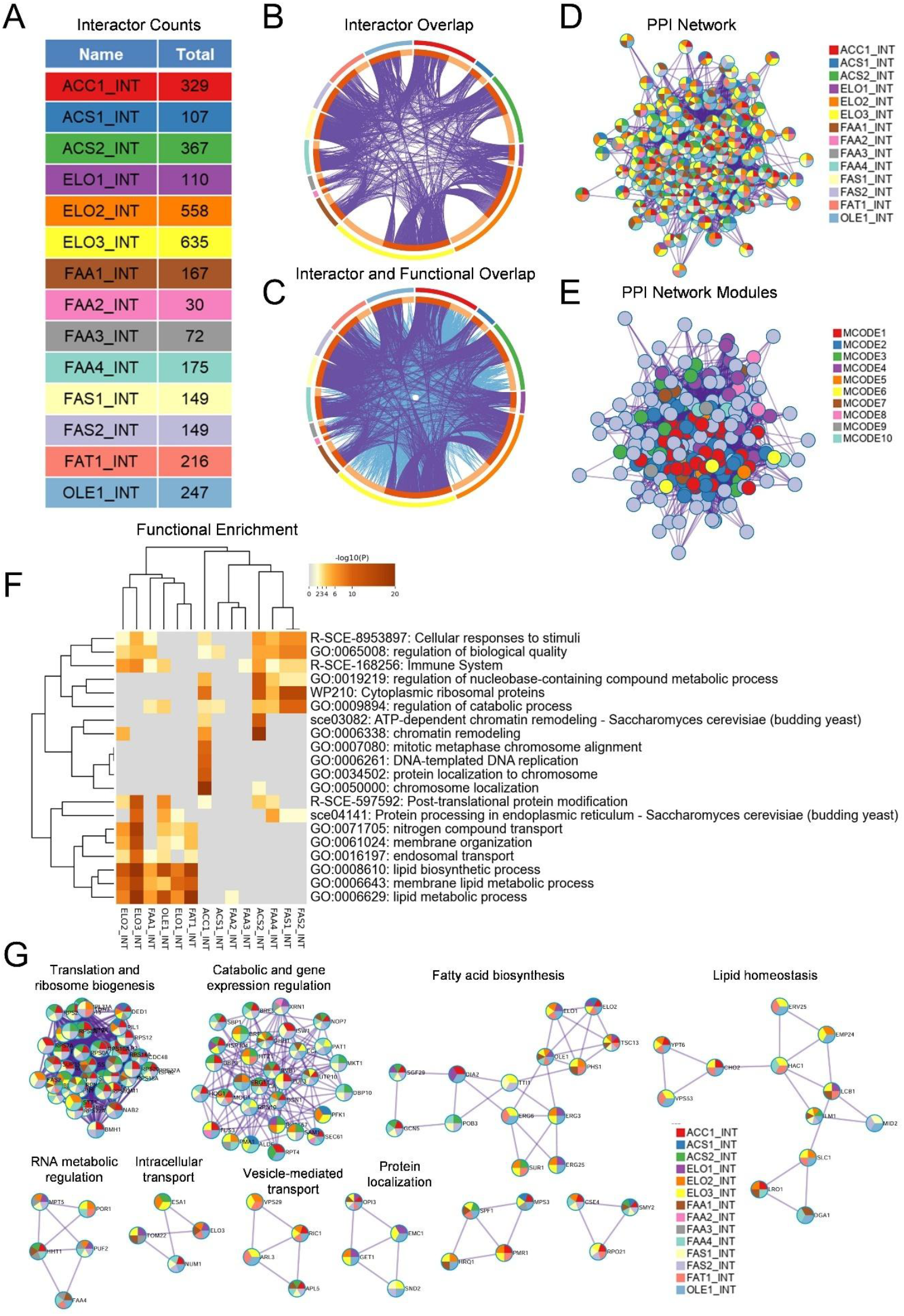
Integrated interactome analysis of yeast fatty acid metabolic pathways. (A) Total physical and genetic interactors identified for yeast fatty acid metabolic regulators, including *ACC1, FAS1, FAS2, ACS1, ACS2, FAA1-4, ELO1-3, OLE1*, and *FAT1* retrieved from the *Saccharomyces* Genome Database (SGD). Each fatty acid metabolic regulator is represented by a distinct color code maintained throughout the figure. (B) Circos plot showing overlap between molecular interactors among yeast fatty acid metabolic regulators. Outer color annotations correspond to individual regulators as shown in panel A. (C) Integrated interactome and functional overlap network illustrating shared molecular interactors and associated biological pathways among yeast fatty acid metabolic regulators. (D) Protein–protein interaction (PPI) network generated from integrated yeast fatty acid metabolic interactors. Node colors indicate corresponding fatty acid metabolic regulators. (E) MCODE-based protein–protein interaction (PPI) network module analysis of the integrated yeast fatty acid metabolic interactome identifying enriched subnetworks. Representative MCODE subnetworks are shown in panel G. (F) Heatmap showing functional enrichment analysis of Gene Ontology (GO) biological processes and pathway terms associated with individual yeast fatty acid metabolic interactomes. Color intensity represents enrichment significance (-log10 P value). (G) Representative MCODE-based protein–protein interaction (PPI) network subnetworks identified from the integrated *ACC1, FAS1, FAS2, ACS1, ACS2, FAA1-4, ELO1-3, OLE1*, and *FAT1* interactome, associated with translation and ribosome biogenesis, catabolic and gene expression regulation, fatty acid biosynthesis, lipid homeostasis, RNA metabolic regulation, intracellular transport, vesicle-mediated transport, and protein localization pathways. Node colors indicate individual fatty acid metabolic interactors according to the corresponding color scheme for each regulator.

Interactor overlap analysis demonstrated extensive sharing of physical and genetic interactors across multiple fatty acid metabolic regulators, indicating strong connectivity between lipid biosynthesis, elongation, activation, desaturation, and membrane-associated metabolic pathways **(Figure 3B)**. Integration of both interactor and functional ontology relationships further revealed broader overlap among fatty acid metabolic pathways and associated biological functions **(Figure 3C)**.

Protein–protein interaction (PPI) network analysis identified densely connected interaction networks among fatty acid metabolic regulators **(Figure 3D)**. Network module analysis further revealed multiple functionally enriched subnetworks associated with translation and ribosome biogenesis, gene expression regulation, fatty acid biosynthesis, lipid homeostasis, RNA metabolism, intracellular transport, vesicle-mediated transport, and protein localization **(Figures 3E and 3G)**.

Functional enrichment analysis revealed two major functional groups within the broader fatty acid metabolic interactome **(Figure 3F)**. One cluster was associated with anabolic growth-related pathways, including cytoplasmic ribosomal proteins, nucleotide metabolism, chromatin remodeling, DNA replication, regulation of gene expression, and catabolic processes. This cluster was predominantly associated with core fatty acid synthesis genes including *FAS1, FAS2, ACS1*, and *ACS2*. Interestingly, *FAA4* also preferentially clustered within this anabolic and stress-associated group, suggesting a potential role in nutrient-responsive metabolic adaptation beyond canonical fatty acid activation pathways. In contrast, a second cluster was enriched for lipid and membrane-associated pathways, including lipid biosynthesis, membrane lipid metabolism, membrane organization, protein processing in the endoplasmic reticulum, endosomal transport, and intracellular transport. Genes involved in fatty acid elongation, desaturation, and membrane lipid remodeling, including *ELO1-3, OLE1, FAA1*, and *FAT1*, preferentially clustered within these membrane-associated pathways. Notably, *ACC1* occupied a more central position between these functional groups, consistent with its role as a key metabolic node linking anabolic growth pathways with downstream fatty acid and membrane lipid biosynthesis. In contrast, *FAA2* and *FAA3* displayed comparatively weaker or more selective ontology enrichment patterns, suggesting functional divergence and context-dependent roles among fatty acid activation pathway components.

Collectively, these findings indicate that yeast fatty acid metabolic pathways are functionally integrated with both anabolic growth regulation and membrane-associated cellular organization, supporting broader roles for lipid metabolic networks in nutrient-responsive growth and stress-regulatory cellular programs.

### Cerulenin-mediated inhibition of fatty acid synthesis suppresses anabolic growth and promotes cellular longevity

Given that the *FAS1/FAS2* and broader fatty acid metabolic interactomes were strongly associated with anabolic growth-related pathways, including translation, ribosome biogenesis, chromatin organization, and gene expression regulation **(Figures 2 and 3)**, we next investigated whether pharmacological inhibition of fatty acid synthesis influences cellular growth and chronological longevity in *Saccharomyces cerevisiae*. For these studies, we used cerulenin, a well-characterized inhibitor of the fatty acid synthase complex targeting *FAS1/FAS2* activity ^37,38^.

Treatment of yeast cells with cerulenin resulted in dose-dependent growth inhibition at both 16 h and 24 h time points **(Figures 4A and 4B)**. Lower concentrations had minimal effects on proliferation, whereas higher concentrations markedly suppressed cellular growth, consistent with inhibition of fatty acid biosynthesis required for anabolic growth and proliferation.

**Figure 4.**
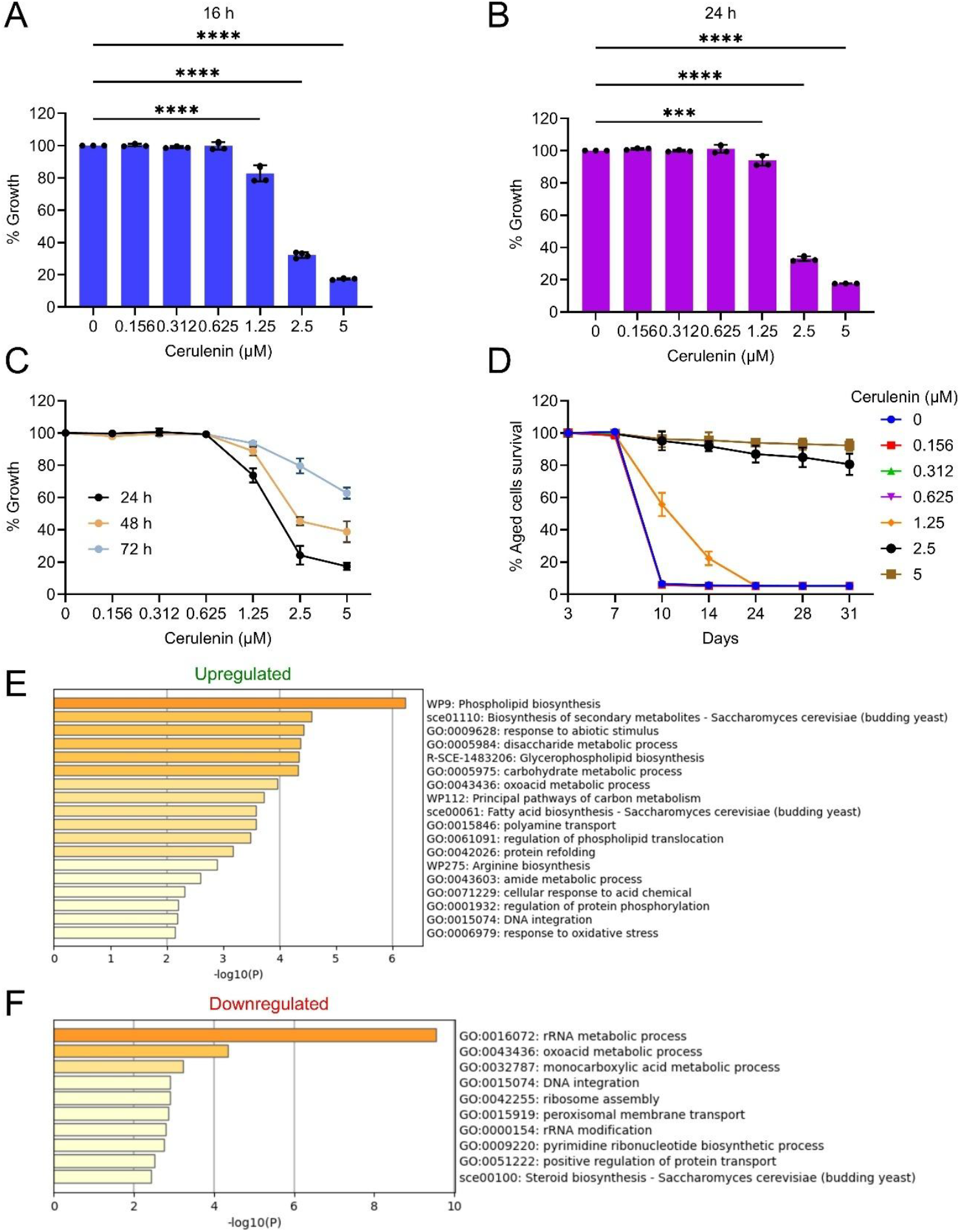
Effects of cerulenin treatment on yeast growth, chronological lifespan, and transcriptomic remodeling. (A,B) Dose-dependent effects of cerulenin on yeast cellular growth following 16 h (A) and 24 h (B) treatment in synthetic defined (SD) medium. Cellular growth was quantified by measuring optical density at 600 nm (OD600). Data are presented as mean ± SD. Statistical significance was determined using one-way ANOVA followed by Dunnett’s multiple comparisons test. ***P < 0.001, ****P < 0.0001. (C) Growth kinetics of yeast cells cultured in the presence of increasing concentrations of cerulenin during transition from exponential growth to stationary phase at 24, 48, and 72 h. Data are presented as mean ± SD (n = 4). (D) Chronological lifespan (CLS) analysis of yeast cells treated with increasing concentrations of cerulenin. Cellular survival during chronological aging was assessed by outgrowth-based viability assays and normalized to day 3 survival. (E,F) Functional enrichment analysis of upregulated (E) and downregulated (F) pathways identified by RNA-sequencing analysis following cerulenin treatment. Color intensity and bar length represent enrichment significance (-log10 P value).

To further examine the effects of cerulenin during transition from exponential growth to stationary phase, cells were cultured in the presence of different cerulenin concentrations and monitored over 24–72 h **(Figure 4C)**. Although higher cerulenin concentrations strongly suppressed growth at early time-points, partial recovery of cellular growth was observed at later stages, particularly during progression toward stationary phase. Since suppression of anabolic and translation-associated pathways is frequently associated with lifespan extension ^39^, we next investigated whether cerulenin treatment influences chronological aging-associated survival.

For chronological lifespan (CLS) analysis, cells were cultured in the presence of cerulenin until post-mitotic stationary phase, with survival measurements initiated after 72 h using outgrowth-based viability assays ^39–43^. Cerulenin treatment markedly extended chronological survival in a dose-dependent manner, with intermediate and higher concentrations maintaining substantially improved long-term survival compared with untreated controls **(Figure 4D)**. These findings indicate that inhibition of fatty acid synthesis promotes enhanced cellular longevity during chronological aging.

To investigate the molecular basis underlying cerulenin-mediated longevity, we performed transcriptomic analysis of exponentially growing cells treated with cerulenin for 1 h to capture early transient transcriptional responses associated with acute fatty acid synthase inhibition. Pathway enrichment analysis revealed upregulation of lipid biosynthesis, phospholipid metabolism, oxidative stress, and arginine biosynthesis-associated pathways, including induction of fatty acid biosynthetic programs **(Figure 4E; Supplementary Data 3)**, suggesting compensatory metabolic responses following fatty acid synthase inhibition. In contrast, cerulenin treatment induced marked downregulation of pathways associated with rRNA metabolism, ribosome assembly, RNA modification, nucleotide biosynthesis, and translation-associated anabolic processes **(Figure 4F; Supplementary Data 3)**.

Overall, these findings indicate that cerulenin-mediated inhibition of fatty acid synthesis suppresses anabolic and translation-associated cellular programs while promoting metabolic remodeling and stress-responsive cellular programs linked to enhanced cellular longevity.

### Fatty acid synthesis inhibition engages TORC1-associated growth regulatory pathways

To further investigate the broader cellular pathways required for adaptation to fatty acid synthase inhibition, we analyzed genome-wide cerulenin-sensitive genetic interaction profiles retrieved from the *Saccharomyces* Genome Database (SGD) and performed functional enrichment analysis **(Figures S1A and S1B; Extended Data 4)**. This analysis identified significant enrichment of pathways associated with fatty-acyl-CoA metabolism, Target of Rapamycin (TOR) signaling, RNA localization, stress granule assembly, phospholipid biosynthesis and broader metabolic regulatory processes, indicating that cellular adaptation to fatty acid synthase inhibition involves coordinated remodeling of anabolic growth and membrane-associated pathways.

Recent studies demonstrated that fatty acid synthase inhibition functionally suppresses Target of Rapamycin Complex 1 (TORC1) signaling through malonyl-CoA-mediated metabolic regulation ^44^. In *Saccharomyces cerevisiae*, amino acid availability is transmitted to TORC1 through the Rag-family GTPase Gtr1, which is anchored to the vacuolar membrane by the Ego complex and regulated by upstream nutrient-sensing pathways ^45^. Consistent with these findings, we next examined whether positive regulators of TORC1 signaling similarly influence cerulenin sensitivity in the prototrophic CEN.PK background and nutrient conditions used in this study. Deletion mutants affecting the TORC1-associated Rag/Gtr signaling pathway, including *GTR1* and the Ego complex component *EGO1*, displayed increased sensitivity to cerulenin treatment compared with wild-type cells **(Figures S1C and S1D)**. Concurrently, these mutants also exhibited increased sensitivity to rapamycin treatment, consistent with impaired TORC1-associated growth regulatory signaling and serving as a positive control for TORC1 pathway dysfunction **(Figures S1E and S1F)**. Together, these findings support conserved functional coupling between fatty acid synthesis and TORC1-associated anabolic growth regulatory pathways across distinct genetic and nutrient contexts.

To further explore broader systems-level relationships associated with fatty acid metabolic regulation, we performed integrative comparative analyses combining fatty acid metabolic regulator interactomes together with cerulenin- and rapamycin-associated transcriptional signatures **(Figures S2A-S2G)**. Comparative clustering analysis revealed segregation between anabolic growth-associated pathways and adaptive metabolic remodeling programs. Cerulenin- and rapamycin-downregulated signatures clustered closely with interactomes associated with core fatty acid biosynthetic regulators, including *FAS1, FAS2, ACS1, ACS2*, and related anabolic-associated pathways linked to translation, ribosome biogenesis, RNA processing, chromatin organization, and broader growth regulatory programs. In contrast, cerulenin- and rapamycin-upregulated signatures preferentially clustered with interactomes associated with fatty acid elongation, desaturation, and membrane-associated regulators, including *ELO1-3, OLE1*, and *FAA*-family proteins, together with pathways linked to lipid metabolism, membrane organization, carbon metabolism, and broader small-molecule metabolic processes. These findings suggest that perturbation of fatty acid synthesis induces coordinated metabolic state transitions involving suppression of anabolic growth-associated programs together with activation of adaptive membrane- and stress-associated metabolic remodeling pathways that partially converge with rapamycin-associated transcriptional responses.

### ER-associated lipid and stress pathways are linked to cellular longevity during chronological aging

Given that lipid biosynthesis primarily occurs at the endoplasmic reticulum (ER) ^20,21^, and that our interactome and transcriptomic analyses consistently identified associations with ER biology, membrane organization, vesicle trafficking, and stress-responsive pathways, we next investigated the relationship between ER-associated lipid metabolic regulation and cellular longevity.

To explore ER-associated interactome organization, we first analyzed the human *DPAGT1* interactome **(Supplementary Data 5)**. *DPAGT1* is a key ER-associated enzyme involved in dolichol-linked glycosylation and protein processing. Functional enrichment analysis revealed significant associations with membrane lipid biosynthetic processes, regulation of protein stability, and regulation of secretion, consistent with central roles in ER-associated membrane and proteostatic organization **(Figure 5A)**.

**Figure 5.**
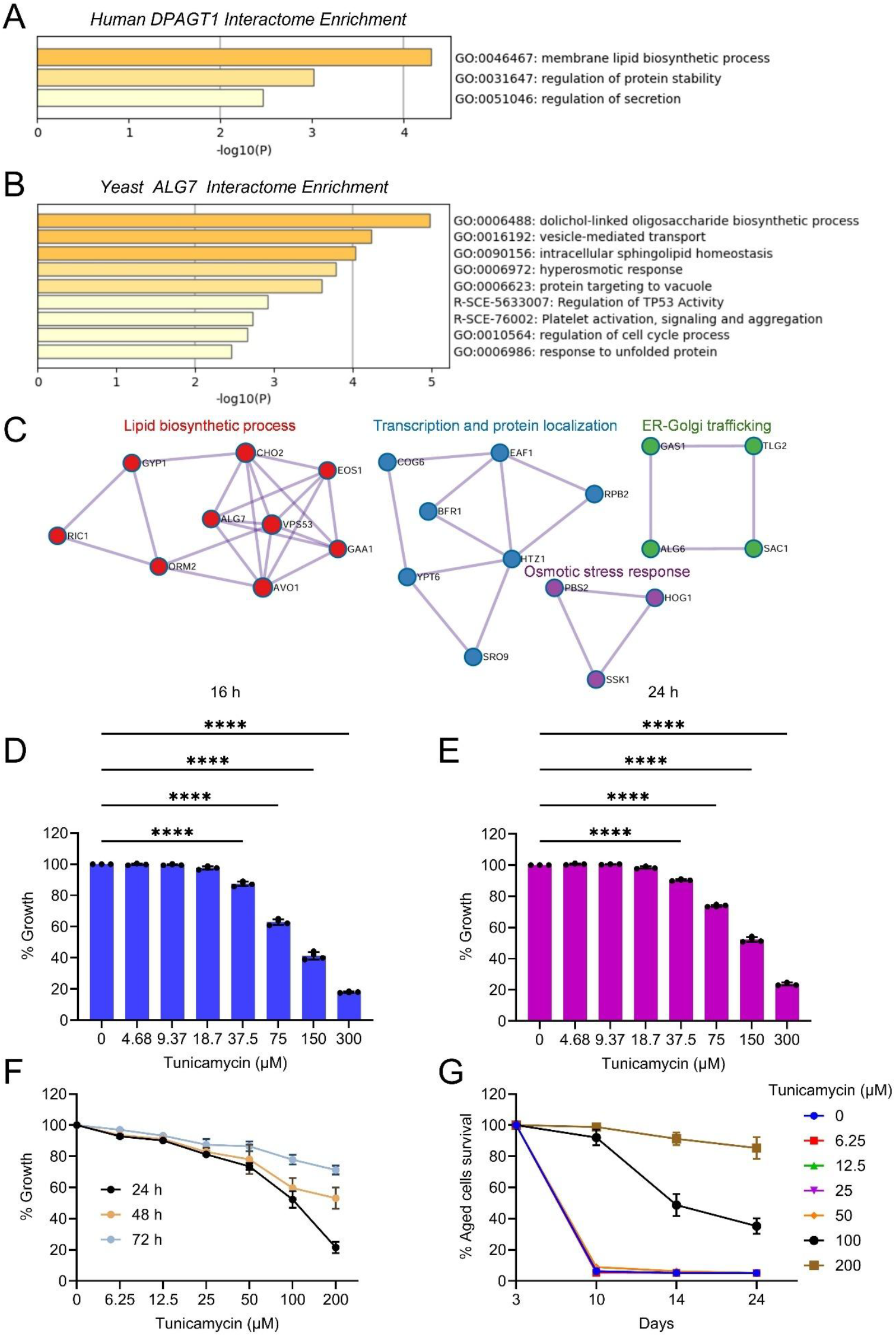
Effects of ER stress modulation on yeast growth and chronological lifespan. (A) Functional enrichment analysis of the human *DPAGT1* interactome. (B) Functional enrichment analysis of the yeast *ALG7* interactome. (C) Representative MCODE-based protein–protein interaction (PPI) network subnetworks identified from the *ALG7* interactome associated with lipid biosynthesis, transcription and protein localization, ER-Golgi trafficking, and osmotic stress response pathways. (D,E) Dose-dependent effects of tunicamycin on yeast cellular growth following 16 h (D) and 24 h (E) treatment in synthetic defined (SD) medium. Cellular growth was quantified by measuring optical density at 600 nm (OD600). Data are presented as mean ± SD. Statistical significance was determined using one-way ANOVA followed by Dunnett’s multiple comparisons test. ****P < 0.0001. (F) Growth kinetics of yeast cells cultured in the presence of increasing concentrations of tunicamycin during transition from exponential growth to stationary phase at 24, 48, and 72 h. Data are presented as mean ± SD (n = 4). (G) Chronological lifespan (CLS) analysis of yeast cells treated with increasing concentrations of tunicamycin. Cellular survival during chronological aging was assessed by outgrowth-based viability assays and normalized to day 3 survival.

To further investigate conserved ER-associated pathways in yeast, we next analyzed the interactome of *ALG7*, the yeast ortholog of *DPAGT1* **(Supplementary Data 5)**. Functional enrichment analysis identified strong associations with dolichol-linked oligosaccharide biosynthetic processes, vesicle-mediated transport, intracellular sphingolipid homeostasis, osmotic stress responses, protein targeting pathways, unfolded protein responses, and regulation of cell cycle-associated processes **(Figure 5B)**. Network module analysis further identified functional subnetworks associated with lipid biosynthesis, transcription and protein localization, ER-Golgi trafficking, and osmotic stress signaling pathways involving the HOG pathway regulator *HOG1* **(Figure 5C)**.

We next examined whether pharmacological ER stress modulation influences cellular growth and chronological longevity using tunicamycin, an inhibitor of N-linked glycosylation that induces ER stress ^46^. Treatment with tunicamycin resulted in dose-dependent suppression of cellular growth at both 16 h and 24 h time points **(Figures 5D and 5E)**, consistent with disruption of ER-associated protein processing and membrane homeostasis. Growth monitoring over 24–72 h further demonstrated progressive growth inhibition at higher tunicamycin concentrations during transition into stationary phase **(Figure 5F)**.

Since cerulenin-mediated longevity was associated with ER, membrane, and stress-associated pathways, we next investigated whether ER stress modulation itself influences chronological aging-associated survival. Chronological lifespan analysis demonstrated that tunicamycin treatment markedly extended long-term cellular survival in a dose-dependent manner, with intermediate and higher concentrations maintaining enhanced survival during chronological aging compared with untreated controls **(Figure 5G)**.

Together, these results support a functional link between ER-associated lipid metabolic organization, stress responses, and regulation of cellular longevity during chronological aging.

## DISCUSSION

Fatty acid biosynthesis is traditionally viewed as a central anabolic pathway required for membrane formation and cellular proliferation ^1,2,6,16,17^. However, emerging evidence suggests that lipid metabolic pathways also function as broader regulators of nutrient sensing, membrane homeostasis, proteostasis, and stress adaptation ^4,7,8,10,12,23,27,29^. In the present study, we combined systems-level interactome analyses with functional aging assays to establish an integrated framework linking fatty acid metabolic organization with anabolic growth control, ER-associated stress pathways, and cellular longevity.

A major outcome of this study is the generation of a human fatty acid metabolic interactome atlas integrating fatty acid synthesis, elongation, desaturation, and transport pathways. Our analyses revealed extensive connectivity between fatty acid metabolic regulators and pathways associated with ER organization, membrane trafficking, autophagy, protein stability, and stress-responsive signaling. These findings indicate that fatty acid metabolic pathways are functionally integrated with broader cellular adaptation programs rather than acting solely as isolated lipid biosynthetic modules. Importantly, many of these organizational features were conserved in yeast, including associations with membrane organization, proteostasis, translation-associated anabolic pathways, and ER stress-related signaling, highlighting evolutionary conservation of fatty acid metabolic network organization across eukaryotic systems.

Expansion of the yeast fatty acid metabolic interactome further revealed segregation into two major functional groups: one associated with anabolic growth-related processes including translation, ribosome biogenesis, nucleotide metabolism, and chromatin organization, and another linked to membrane organization, lipid homeostasis, intracellular transport, and ER-associated pathways. Notably, *ACC1* occupied a central position between these clusters, supporting its role as a metabolic node linking anabolic growth programs with downstream fatty acid and membrane lipid biosynthesis. These findings further support emerging concepts proposing that lipid metabolic pathways coordinate broader growth-regulatory and stress-associated cellular programs ^10,15^.

Using yeast chronological lifespan as a tractable aging model, we further demonstrated that pharmacological inhibition of fatty acid synthesis using cerulenin suppresses cellular proliferation while promoting enhanced long-term survival. Transcriptomic analysis revealed broad downregulation of translation-associated and anabolic pathways together with induction of stress adaptation and membrane-associated metabolic programs. One potential mechanistic explanation may involve accumulation of malonyl-CoA following fatty acid synthase inhibition. Recent work demonstrated that malonyl-CoA functions as a conserved endogenous ATP-competitive inhibitor of TORC1 in both yeast and mammalian cells, thereby coupling fatty acid biosynthetic capacity to global anabolic regulation ^44,47^. Consistent with this model, cerulenin treatment in our study strongly suppressed pathways associated with translation, ribosome biogenesis, RNA metabolism, and nucleotide biosynthesis linked to TORC1-dependent anabolic signaling. Moreover, mutants affecting positive regulators of TORC1 signaling, including *GTR1* and the Ego complex component *EGO1*, displayed increased sensitivity to cerulenin treatment in the prototrophic CEN.PK background, further supporting conserved functional coupling between fatty acid synthesis and TORC1-associated anabolic growth regulation under our experimental conditions.

Another important finding from our study is the recurrent association between fatty acid metabolic organization and ER-associated stress pathways. Both *DPAGT1* and *ALG7* interactome analyses identified strong enrichment of protein processing, vesicle trafficking, unfolded protein response-associated pathways, and membrane homeostasis networks.

Moreover, pharmacological induction of ER stress using tunicamycin extended chronological lifespan despite suppressing cellular growth. Increasing evidence suggests that adaptive ER stress and membrane stress signaling pathways contribute to metabolic remodeling and longevity-associated cellular adaptation ^48–52^. Importantly, mild ER stress responses have previously been shown to promote longevity in *Caenorhabditis elegans*, suggesting evolutionary conservation of adaptive ER-associated stress mechanisms across species ^50,53^.

Collectively, our integrated fatty acid metabolic interactome atlases uncovered conserved biological signatures linking anabolic growth regulation, membrane organization, proteostasis, and ER-associated stress pathways across human and yeast systems. Functional interrogation of these conserved network signatures demonstrated that pharmacological inhibition of fatty acid synthesis using cerulenin promotes adaptive metabolic remodeling associated with enhanced longevity in yeast. Moreover, pharmacological induction of ER stress using tunicamycin, previously implicated in longevity-associated adaptation in other model systems, similarly promoted enhanced cellular survival during chronological aging. More broadly, this study demonstrates how integrated fatty acid metabolic interactome atlases combined with transcriptomic and functional aging approaches can identify conserved lipid metabolic mechanisms linked to aging biology and uncover candidate longevity-associated interventions.

## METHODS

### Human and yeast fatty acid metabolic interactome analysis

Molecular interactors of human fatty acid metabolic regulators involved in fatty acid synthesis, activation, elongation, desaturation, and transport, including *ACACA, FASN, ACSS1, ACSS2, ACSL1, ACSL3, ACSL4, ACSL5, ACSL6, ELOVL1, ELOVL2, ELOVL3, ELOVL4, ELOVL5, ELOVL6, ELOVL7, SCD, SCD5, SLC27A1, SLC27A2, SLC27A3, SLC27A4, SLC27A5*, and *SLC27A6*, were retrieved from the Alliance of Genome Resources database. Physical and genetic interactors of the yeast orthologs *ACC1, FAS1, FAS2, ACS1, ACS2, FAA1, FAA2, FAA3, FAA4, ELO1, ELO2, ELO3, OLE1*, and *FAT1* were obtained from the *Saccharomyces* Genome Database (SGD). Integrated interactome and functional enrichment analyses were subsequently performed using the Metascape online platform, which integrates multiple bioinformatics resources for pathway enrichment and protein–protein interaction (PPI) network analysis ^36^. Default analysis parameters were used for all analyses. Significantly enriched Gene Ontology (GO) biological processes, pathway terms, interactome overlap networks, and PPI network modules were visualized to identify conserved biological signatures and functional pathways associated with fatty acid metabolic interactomes.

### Yeast strains and culture conditions

The prototrophic *Saccharomyces cerevisiae* CEN.PK113-7D wild-type strain was used throughout this study ^76^. Gene deletion mutants were generated using PCR-mediated homologous recombination as previously described ^54^. Yeast cells were recovered from frozen glycerol stocks on YPD agar plates containing 1% yeast extract, 2% peptone, 2% glucose, and 2% agar, followed by incubation at 30 °C for 2–3 days. For experimental analyses, cells were cultured in synthetic defined (SD) medium containing 6.7 g/L yeast nitrogen base with ammonium sulfate (without amino acids) supplemented with 2% glucose.

### Chemical treatments

Cerulenin and tunicamycin were prepared as stock solutions in dimethyl sulfoxide (DMSO) and added to yeast cultures at the indicated concentrations. Control cultures received equivalent volumes of vehicle alone. The final concentration of DMSO did not exceed 1% in any experiment.

### Growth assay

Yeast growth assays were performed in a 96-well microplate format to evaluate the effects of chemical treatments on cellular proliferation. Prototrophic *Saccharomyces cerevisiae* CEN.PK113-7D wild-type and the mutant strains were cultured in synthetic defined (SD) medium containing 6.7 g/L yeast nitrogen base with ammonium sulfate (without amino acids) supplemented with 2% glucose. Cells were inoculated at an initial optical density of approximately OD600 = 0.2 and dispensed into 96-well plates containing 200 μL culture volume per well with serial two-fold dilutions of the indicated compounds. Plates were incubated at 30 °C, and cellular growth was monitored by measuring optical density at 600 nm (OD600) using a microplate reader.

### Chronological lifespan assay

Chronological lifespan (CLS) assays were performed as described previously ^39–41^. *Saccharomyces cerevisiae* prototrophic CEN.PK113-7D cells were cultured in synthetic defined (SD) medium containing 6.7 g/L yeast nitrogen base with ammonium sulfate (without amino acids) supplemented with 2% glucose. Overnight cultures grown at 30 °C with shaking at 220 rpm were diluted to an initial optical density of approximately OD600 = 0.2 in fresh SD medium and dispensed into 96-well plates containing 200 μL culture volume per well to initiate CLS experiments. Cultures were maintained at 30 °C, and growth was monitored during transition from exponential growth to stationary phase at 24, 48, and 72 h. At the indicated chronological aging time points, 2 μL of aged stationary-phase culture was transferred into a second 96-well plate containing 200 μL YPD medium and incubated for 24 h at 30 °C. Cellular outgrowth was quantified by measuring optical density at 600 nm (OD600) using a microplate reader. The 72 h (day 3) time point was designated as 100% survival and used as the reference point for normalization of survival measurements at subsequent chronological aging time points.

### RNA extraction

*Saccharomyces* cerevisiae prototrophic CEN.PK113-7D cells were cultured in synthetic defined (SD) medium containing 6.7 g/L yeast nitrogen base with ammonium sulfate (without amino acids) supplemented with 2% glucose. Exponentially growing cultures were treated with cerulenin (5 μM) for 1 h prior to cell harvesting to capture early transcriptional responses to fatty acid synthase inhibition. Yeast cells were mechanically lysed according to the manufacturer’s recommended protocol, and total RNA was extracted using the RNeasy Mini Kit (Qiagen). RNA concentration and purity were assessed using an ND-1000 UV–visible spectrophotometer (NanoDrop Technologies) ^55,56^. High-quality RNA samples were subsequently used for RNA-sequencing analysis as described below.

### RNA sequencing and functional enrichment analysis

RNA integrity was evaluated using the Agilent 2100 Bioanalyzer with the RNA 6000 Nano LabChip kit. High-quality RNA samples were subjected to paired-end RNA sequencing at Novogene. Polyadenylated mRNA was enriched prior to library construction, followed by cDNA synthesis through reverse transcription. Library quality and fragment size distribution were verified using the Bioanalyzer, and sequencing was performed on the NovaSeq PE150 platform. Raw sequencing reads were processed using the nf-core RNA-seq pipeline (version 3.8.1), employing STAR for read alignment and RSEM for transcript quantification ^57^. Reads were aligned to the Saccharomyces cerevisiae reference genome using corresponding Ensembl gene annotations (GTF version 1.105), generating gene-level expression counts ^58^. Normalization and statistical analysis of gene expression were performed using DESeq2 ^59^. For pathway-level analysis of early transcriptional responses, genes with nominal P values < 0.05 were selected. Genes with positive log2 fold-change values were classified as upregulated, whereas genes with negative log2 fold-change values were classified as downregulated. Rapamycin-associated RNA-sequencing datasets used in this study were obtained from our previous work ^55^. Functional annotation and enrichment analyses of these nominally responsive gene sets were performed using the Metascape online platform ^36^.

### Statistical analysis

All statistical analyses and data visualization were performed using GraphPad Prism (version 11). Data are presented as mean ± standard deviation (SD) unless otherwise indicated. Statistical significance was determined using one-way analysis of variance (ANOVA), with multiple-comparison corrections applied as specified in the corresponding figure legends.

## Code availability

No custom code or mathematical algorithm was used in this study.

## Data availability

Further information and requests for resources and reagents should be directed to and will be fulfilled by the Lead Contact, Dr. Mohammad Alfatah.

## ACKNOWLEDGMENTS

This work is supported by the Young Investigator Research Grant (YIRG), National Medical Research Council, Singapore (MOH-001348-00) and US NAM Healthy Longevity Catalyst Awards Grant (MOH-001439). Arshia Naaz is supported by A*STAR-CDF grant (C243512027).

## AUTHOR CONTRIBUTIONS STATEMENT

Arshia Naaz: Writing-original draft, Investigation, Formal analysis, Funding Acquisition

Mingtong Gao: Investigation, Formal analysis

Yizhong Zhnag: Investigation, Formal analysis

Rajkumar Dorajoo: Review and editing

Brian K Kennedy: Review and editing

Mohammad Alfatah: Conceptualization, Writing-review and editing, Funding acquisition.

All authors read, critically reviewed and approved the final manuscript.

M.A is the guarantor of this work.

