## Supplemental Figures for "Fatty acid metabolic interactome atlas linked to cellular longevity"

Affiliations:
1. Genome Institute of Singapore (GIS), Agency for Science, Technology and Research (A*STAR), 60 Biopolis Street, Genome #02-01, Singapore 138672, Republic of Singapore.
2. Healthy Longevity Translational Research Program, Yong Loo Lin School of Medicine, National University of Singapore, Singapore, Singapore.
3. Centre for Healthy Longevity, National University Health System, Singapore, Singapore.
4. Bioinformatics Institute (BII), Agency for Science, Technology and Research (A*STAR), 30 Biopolis Street, Matrix #07-01, Singapore 138671, Republic of Singapore.
5. Department of Paediatrics, Yong Loo Lin School of Medicine, National University of Singapore, Singapore 119228, Singapore.
6. Departments of Biochemistry and Physiology, Yong Loo Lin School of Medicine, National University of Singapore, Singapore, Singapore.

*To whom the correspondence should be addressed.

 (Mohammad Alfatah)

 (Arshia Naaz)

**Keywords:** Interactome; Fatty acid metabolism; Lipid metabolism; Endoplasmic reticulum stress; Aging; Cellular longevity

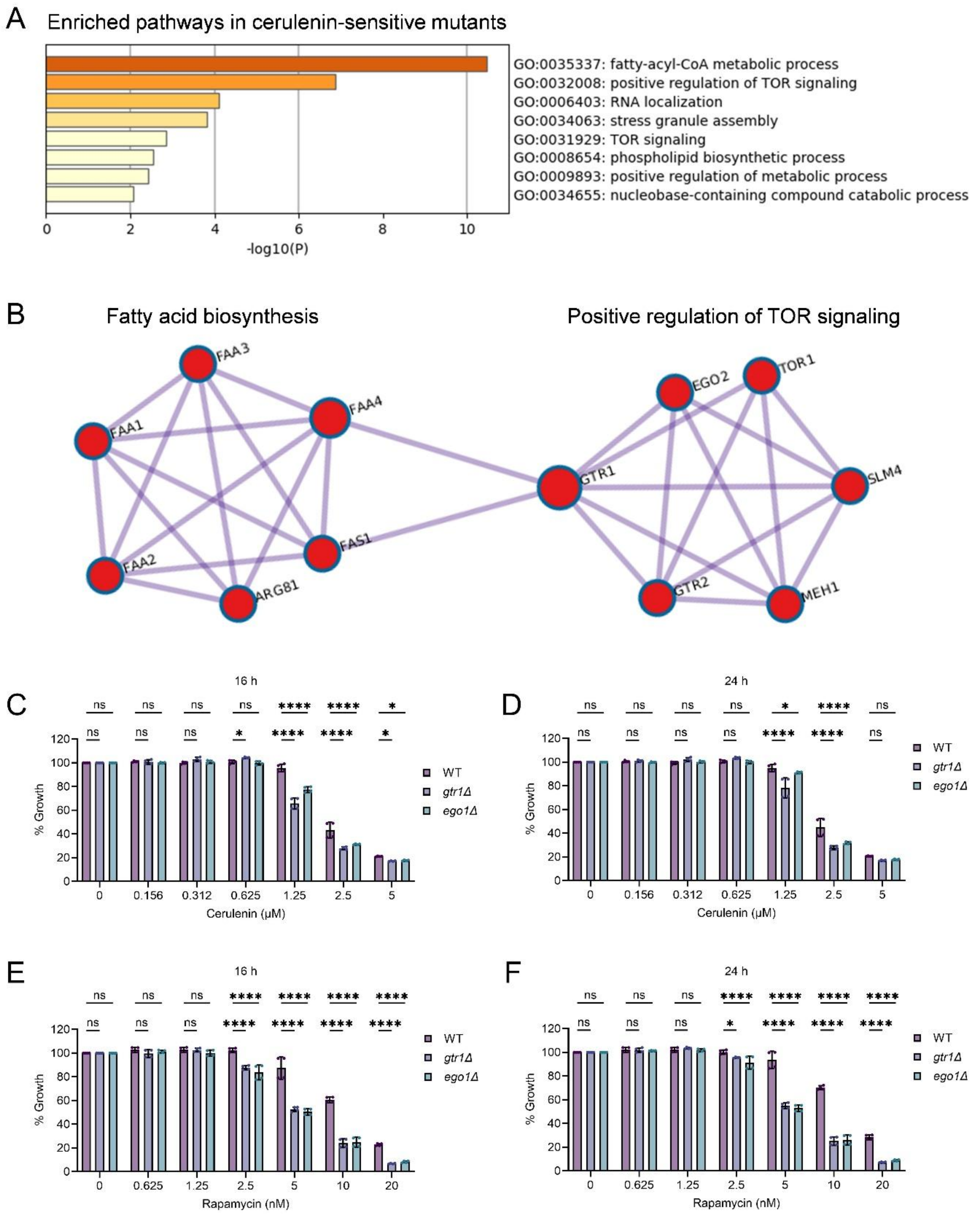


**Figure S1. Fatty acid synthesis inhibition engages Target of Rapamycin Complex 1 (TORC1)-associated growth regulatory pathways**

(A) Functional enrichment analysis of pathways associated with cerulenin-sensitive mutants retrieved from the Saccharomyces Genome Database (SGD). Bar length and color intensity represent enrichment significance (-log10 P value).

(B) MCODE-based protein–protein interaction (PPI) network analysis identifying interconnected subnetworks associated with fatty acid biosynthesis and positive regulation of TOR signaling pathways among cerulenin-sensitive mutants.

(C,D) Growth analysis of wild-type (WT), *gtr1Δ*, and *ego1Δ* mutant strains cultured in the presence of increasing concentrations of cerulenin for 16 h (C) and 24 h (D). Data are presented as mean ± SD ($n = 4$). Statistical significance was determined using two-way ANOVA with Dunnett's multiple comparisons test comparing mutant strains with WT controls. ns, not significant; *$P < 0.05$; ****$P < 0.0001$.

(E,F) Growth analysis of WT, *gtr1Δ*, and *ego1Δ* mutant strains cultured in the presence of increasing concentrations of rapamycin for 16 h (E) and 24 h (F). Data are presented as mean ± SD ($n = 4$). Statistical significance was determined using two-way ANOVA with Dunnett's multiple comparisons test comparing mutant strains with WT controls. ns, not significant; *$P < 0.05$; ****$P < 0.0001$.

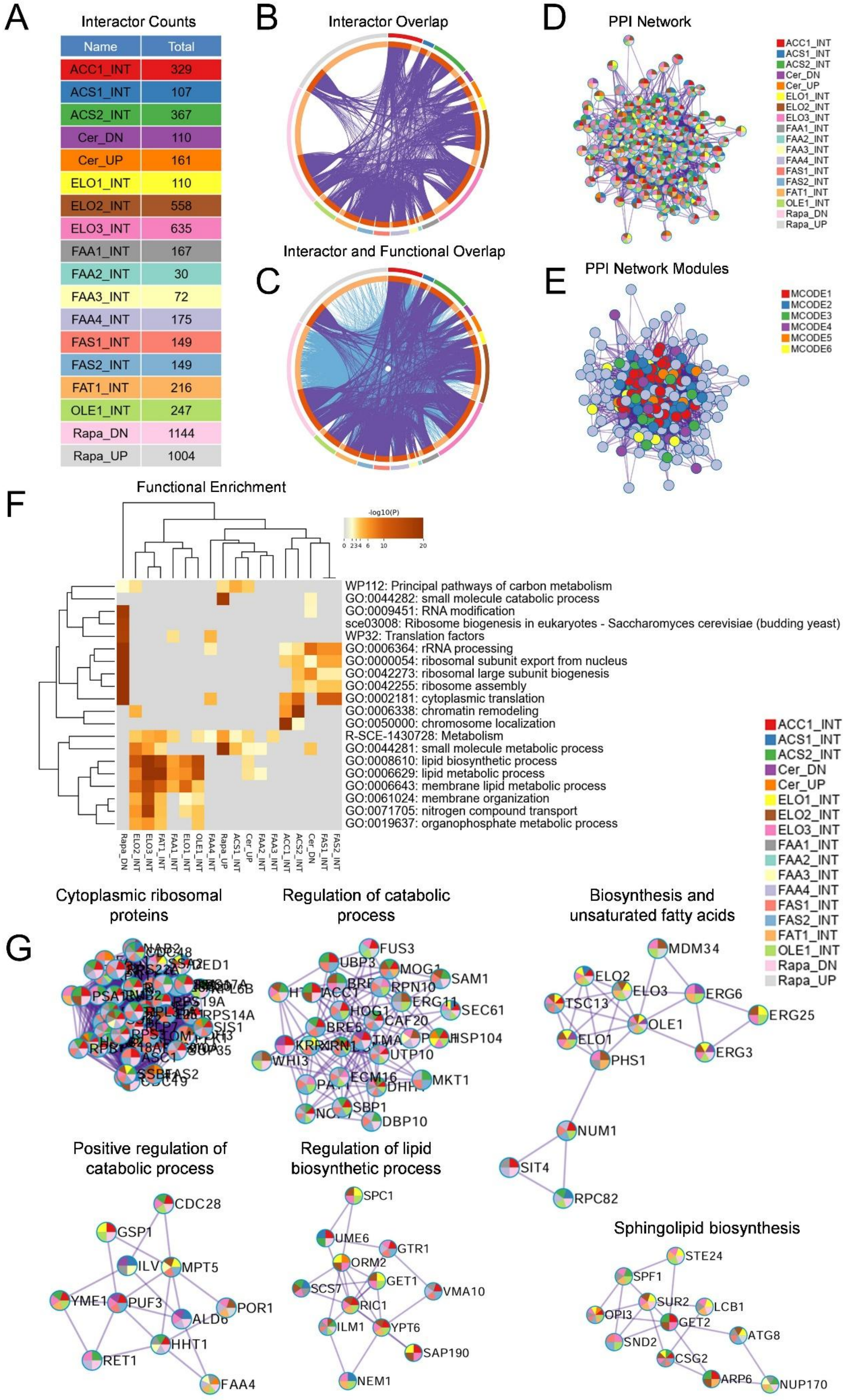
A
Interactor Counts
Name Total
ACC1_INT 329
ACS1_INT 107
ACS2_INT 367
Cer_DN 110
Cer_UP 161
ELO1_INT 110
ELO2_INT 558
ELO3_INT 635
FAA1_INT 167
FAA2_INT 30
FAA3_INT 72
FAA4_INT 175
FAS1_INT 149
FAS2_INT 149
FAT1_INT 216
OLE1_INT 247
Rapa_DN 1144
Rapa_UP 1004
B
Interactor Overlap
C
Interactor and Functional Overlap
D
PPI Network
ACC1_INT
ACS1_INT
ACS2_INT
Cer_DN
Cer_UP
ELO1_INT
ELO2_INT
ELO3_INT
FAA1_INT
FAA2_INT
FAA3_INT
FAA4_INT
FAS1_INT
FAS2_INT
FAT1_INT
OLE1_INT
Rapa_DN
Rapa_UP
E
PPI Network Modules
MCODE1
MCODE2
MCODE3
MCODE4
MCODE5
MCODE6
F
Functional Enrichment
-log10(P)
0 2 3 4 6 10 20
WP112: Principal pathways of carbon metabolism
GO:0044282: small molecule catabolic process
GO:0009451: RNA modification
sce03008: Ribosome biogenesis in eukaryotes - Saccharomyces cerevisiae (budding yeast)
WP32: Translation factors
GO:0006364: rRNA processing
GO:0000054: ribosomal subunit export from nucleus
GO:0042273: ribosomal large subunit biogenesis
GO:0042255: ribosome assembly
GO:0002181: cytoplasmic translation
GO:0006338: chromatin remodeling
GO:0050000: chromosome localization
R-SCE-1430728: Metabolism
GO:0044281: small molecule metabolic process
GO:0008610: lipid biosynthetic process
GO:0006629: lipid metabolic process
GO:0006643: membrane lipid metabolic process
GO:0061024: membrane organization
GO:0071705: nitrogen compound transport
GO:0019637: organophosphate metabolic process
Rapa_DN ELO2_INT ELO3_INT FAT1_INT FAA1_INT ELO1_INT OLE1_INT FAA4_INT Rapa_UP ACS1_INT Cer_UP FAA2_INT FAA3_INT ACC1_INT ACS2_INT Cer_DN FAS1_INT FAS2_INT
ACC1_INT
ACS1_INT
ACS2_INT
Cer_DN
Cer_UP
ELO1_INT
ELO2_INT
ELO3_INT
FAA1_INT
FAA2_INT
FAA3_INT
FAA4_INT
FAS1_INT
FAS2_INT
FAT1_INT
OLE1_INT
Rapa_DN
Rapa_UP
G
Cytoplasmic ribosomal proteins
Regulation of catabolic process
FUS3
UBP3
MOG1
SAM1
RPN10
ERG11
SEC61
HOG1
CAF20
BRE5
HSP104
UTP10
WHI3
ECM16
MKT1
SBP1
DBP10
Biosynthesis and unsaturated fatty acids
MDM34
ELO2
ELO3
ERG6
TSC13
OLE1
ERG25
ELO1
ERG3
PHS1
NUM1
SIT4
RPC82
Positive regulation of catabolic process
CDC28
GSP1
MPT5
YME1
PUF3
POR1
HHT1
RET1
FAA4
Regulation of lipid biosynthetic process
SPC1
UME6
GTR1
ORM2
GET1
SCS7
VMA10
RIC1
ILM1
YPT6
SAP190
NEM1
Sphingolipid biosynthesis
STE24
SPF1
SUR2
LCB1
OPI3
GET2
SND2
ATG8
CSG2
ARP6
NUP170

**Figure S2. Integrative comparative analysis of yeast fatty acid metabolic interactomes with cerulenin- and rapamycin-associated transcriptional signatures**

(A) Total physical and genetic interactors identified for yeast fatty acid metabolic regulators, including *ACC1, FAS1, FAS2, ACS1, ACS2, FAA1-4, ELO1-3, OLE1,* and *FAT1* retrieved from the *Saccharomyces* Genome Database (SGD), together with differentially expressed gene signatures identified from cerulenin-treated (Cer_UP and Cer_DN) and rapamycin-treated (Rapa_UP and Rapa_DN) transcriptomic datasets. Each interactome and transcriptional signature is represented by a distinct color code maintained throughout the figure.

(B) Circos plot showing overlap between molecular interactors and transcriptional signatures among yeast fatty acid metabolic regulators together with cerulenin- and rapamycin-associated gene expression datasets. Outer color annotations correspond to individual datasets shown in panel A.

(C) Integrated interactome and functional overlap network illustrating shared molecular interactors, transcriptional signatures, and associated biological pathways among yeast fatty acid metabolic regulators together with cerulenin- and rapamycin-responsive genes.

(D) Protein–protein interaction (PPI) network generated from integrated yeast fatty acid metabolic interactors and cerulenin- and rapamycin-associated transcriptional signatures. Node colors indicate corresponding interactomes or transcriptional datasets.

(E) MCODE-based protein–protein interaction (PPI) network module analysis of the integrated fatty acid metabolic interactome and transcriptomic datasets identifying enriched functional subnetworks. Representative MCODE subnetworks are shown in panel G.

(F) Heatmap showing functional enrichment analysis of Gene Ontology (GO) biological processes and pathway terms associated with individual yeast fatty acid metabolic interactomes together with cerulenin- and rapamycin-associated transcriptional signatures. Color intensity represents enrichment significance (-log10 P value).

(G) Representative MCODE-based protein–protein interaction (PPI) network subnetworks identified from the integrated fatty acid metabolic interactome and transcriptomic datasets.
